# High resolution data-driven model of the mouse connectome

**DOI:** 10.1101/293019

**Authors:** Joseph E. Knox, Kameron Decker Harris, Nile Graddis, Jennifer D. Whitesell, Hongkui Zeng, Julie A. Harris, Eric Shea-Brown, Stefan Mihalas

## Abstract

Knowledge of mesoscopic brain connectivity is important for understanding inter- and intra-region information processing. Models of structural connectivity are typically constructed and analyzed with the assumption that regions are homogeneous. We instead use the Allen Mouse Brain Connectivity Atlas to construct a model of whole brain connectivity at the scale of 100 µm voxels. The dataset used consists of 366 anterograde tracing experiments in wild type C7BL/6 mice, mapping fluorescently-labeled neuronal projections brain-wide. Inferring spatial connectivity with this dataset remains underdetermined, since the approximately 2 × 10^5^ source voxels outnumber the number of experiments. To address this, we assume that connection patterns and strengths vary smoothly across major brain divisions. We model the connectivity at each voxel as a radial basis kernel-weighted average of the projection patterns of nearby injections. The voxel model outperforms a previous regional model in predicting held-out experiments and compared to a human-curated dataset. This voxel-scale model of the mouse connectome permits researchers to extend their previous analyses of structural connectivity to unprecedented levels of resolution, and allows for comparison with functional imaging and other datasets.

## 1 Introduction

Brain network structure, across many spatial scales, plays an important role in facilitating and constraining neural computations. Models of structural connectivity have been used to investigate the relationship with functional connectivity, to compare brain structures across species, and more (Laramée & Boire, 2015; Sethi, Zerbi, Wenderoth, Fornito, & Fulcher, 2017; Stafford et al., 2014; Wang & Kennedy, 2016). However, most of our knowledge of neuronal networks is limited to either detailed description of small systems (Bock et al., 2011; Glickfeld, Andermann, Bonin, & Reid, 2013; Kleinfeld et al., 2011; White, Southgate, Thomson, & Brenner, 1986) or to a coarse description of connectivity between larger regions (Felleman & Van Essen, 1991; Sporns, 2010). In between these two extremes is *mesoscopic* structural connectivity: a coarser scale than that of single neurons or cortical columns but finer than whole brain regions. Facilitated by new tracing techniques, image processing algorithms, and high-throughput methods, mesoscale data with full brain coverage exist in animals such as the fly (Jenett et al., 2012; Peng et al., 2014) and mouse (Gămănuţ et al., 2018; Kuan et al., 2015; Oh et al., 2014), and such data are being collected from other model organisms (Bota, Dong, & Swanson, 2003; Majka et al., 2016).

We present a scalable regression technique for constructing spatially explicit mesoscale connectivity from anterograde tracing experiments. Specifically, our model estimates the projection strength between every pair of approximately 5 × 10^5^ cubic voxels, each 100 µm wide, in the mouse brain. We use data from the Allen Mouse Brain Connectivity Atlas (Oh et al., 2014), a large scale dataset viral tract-tracing experiments performed across many regions of the mouse brain. All of the data and scripts are publicly available at https://github.com/AllenInstitute/voxel_model.

In these mesoscale anterograde tracing experiments, a tracer virus (recombinant adeno-associated virus, or rAAV) is first injected into the brain. The virus rAAV infects neurons at the site of injection, and causes them to express green fluorescent protein in their cytoplasm, including throughout the entire length of their axons. Brains and labeled axons are imaged with serial two-photon tomography throughout the entire rostral-to-caudal extent of the brain, resulting in an aligned stack of 2-D images that can easily be transformed to 3-D space. Each brain contains one source injection only. Every image series is registered to the 3D Allen Mouse Brain Reference Atlas space, using a combination of global affine and local transformations (Kuan et al., 2015). The reference atlas provides a common coordinate framework for data integration at the voxel level, and is fully annotated with structure/region delineations.

Combining many experiments with different sources then reveals the pathways that connect those sources throughout the brain, the ingredients of a “connectome”. This requires combining data across multiple animals, which appears justified at the mesoscale (Oh et al., 2014). Previous mouse connectome models were constructed with the assumption that regions are homogeneous (Gămănuţ et al., 2018; Oh et al., 2014). While these have proven useful, they depend on predefined regional parcelations and describe connectivity at a region-limited level of resolution. Here, we go beyond the regional approach and construct a model of the whole brain connectivity at the scale of 100 µm voxels.

Previously, Harris, Mihalas, and Shea-Brown (2016) formulated a regularized, structured regression problem for inferring voxel connectivity. This model was applied to Allen Mouse Brain Connectivity Atlas data in the visual cortex, outperforming a regional model in prediction of held-out experiments. We have now extended this approach from the visual cortex to the full mouse brain, while simplifying the mathematical model for computational efficiency. Our model relaxes the assumption of homogeneity of connections within a region and instead assumes smoothness across major brain divisions. We model the connectivity at each source voxel as the weighted average of the projection patterns of nearby injections, where the weights are a nonlinear function of distance to the injection centroid. We fit the parameters of the model using nested cross-validation with held out injection experiments. The new voxel-scale model generally outpredicts a homogeneous regional model, as measured both by cross-validation error and when compared to a human-curated dataset.

The new voxel-scale model will be useful for many applications. For example, we recently used it to reveal the community structure of the mouse cortical network, where the model provided data for regions lacking well-isolated injection experiments. We believe that this is the tip of the iceberg, and that this new voxel-scale model of the mouse connectome will permit researchers to extend their previous analyses of structural connectivity with unprecedented levels of resolution.

## 2 Results

### 2.1 Spatial method to infer the voxel connectome

Here, we will give a short overview of our method for connectome regression. The details are found in Methods. We consider the problem of fitting a weighted, directed, adjacency matrix which contains the connection strength between any pair of points in the brain. We use *n* cubic voxels, 100 µm across, to discretize the brain volume. Our goal is then to find a matrix 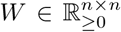 that accurately captures voxel-voxel connection strength. We assume there exists some underlying matrix *W* that is common across animals. Each experiment can be thought of as an injection *X*, and its projections *Y*, where *X*, *Y* ∈ ℝ^*n*^, and we want to find *W* so that *Y* ≈ *WX*, i.e. we want to solve a multivariate regression problem.

We adopt a spatial weighting technique to combine information from multiple experiments into one matrix, the outline of which is shown in Fig. 1. As in Harris et al. (2016), we assume that the connectivity from any given source voxel varies smoothly as a function of distance: columns *W*_:,*i*_ and *W*_:,*j*_ should be similar if the distance between voxels *i* and *j* is small. We make the mathematically simplifying assumption that the projections we observe from a given experiment come from the center of mass of the injection *c*_*e*_. This allows us to employ kernel regression to approximate the connectivity from a given voxel *v* as the distance-weighted sum of injections in the major brain division containing *v*. We also expect the connectivity could change sharply between the boundaries of high-level brain structures. For example, we know that projections from the thalamus and hypothalamus are very different, even though some areas within these major regions are near each other at the borders. To account for this, we partition the brain into 12 distinct major brain divisions. These 12 major brain divisions constitute the set of *coarse structures* defined in the 3-D Allen Mouse Brain Reference Atlas. These areas are: Isocortex, Olfactory Bulb, Hippocampus, Cortical Subplate, Striatum, Pallidum, Thalamus, Hypothalamus, Midbrain, Pons, Medulla, Cerebellum.

**Figure 1:**
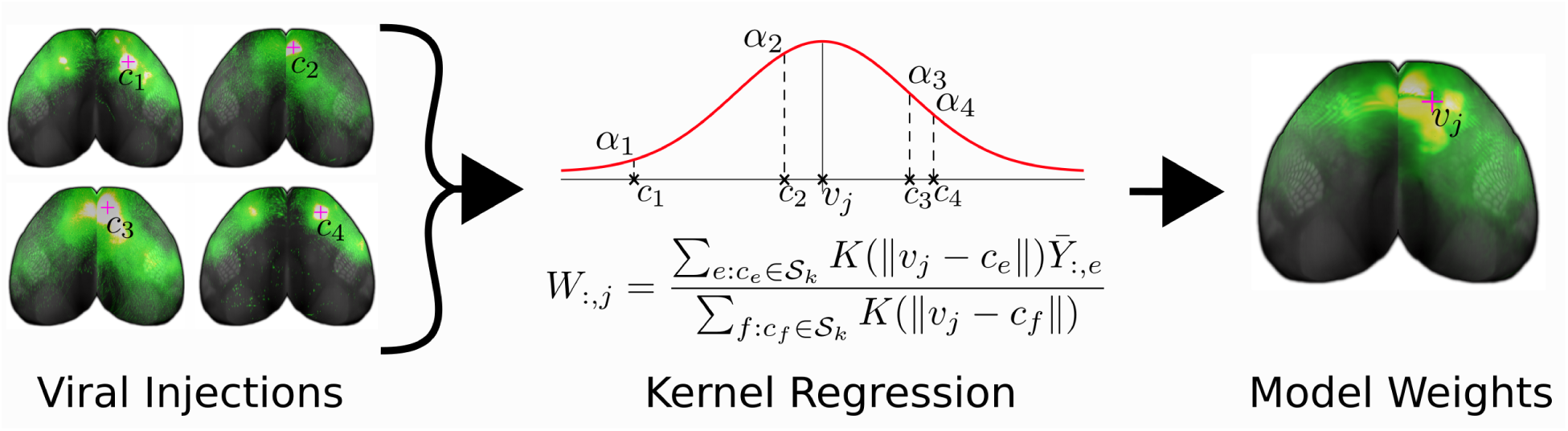
Cartoon illustrating the overview of our method. We combine the information from many viral tracing experiments with different injection sites into a model of voxel structural connectivity. To predict the weight of projections from a voxel *v*, we take an average of nearby injections where the *e*th experiment is weighted by a factor proportional to *K*(||*v* − *c*_*e*_||) and *K*(⋅) is the kernel.

### 2.2 Voxel-scale model outperforms previous regionally homogeneous model

Oh et al. (2014) obtained a regional connectome by integrating the injection and projection data over regions and fitting a region-by-region matrix with nonnegative least squares. We recomputed such a matrix (see Methods) and compared it to a *regionalized* version of the voxel connectivity. To avoid confusion, we call this the regionally *homogeneous* model. In short, we chose 292 regions which are intermediate level “summary structures” in the 3-D Allen Mouse Brain Reference Atlas. We recomputed the *homogeneous* model since the Allen Mouse Brain Reference Atlas was updated in the meantime. The voxel connectivity was then integrated and averaged over regions to produce regionalized weights (see Methods for details).

In Fig. 2, there is a depiction of the whole-brain regionalized weights and, in Fig. 3, the regionalized weights for isocortex (the largest major brain division). Note that, for visualization purposes, we depict sources as rows and targets as columns. This is the opposite of our mathematical convention, so we are in fact depicting *W*^*T*^.

**Figure 2:**
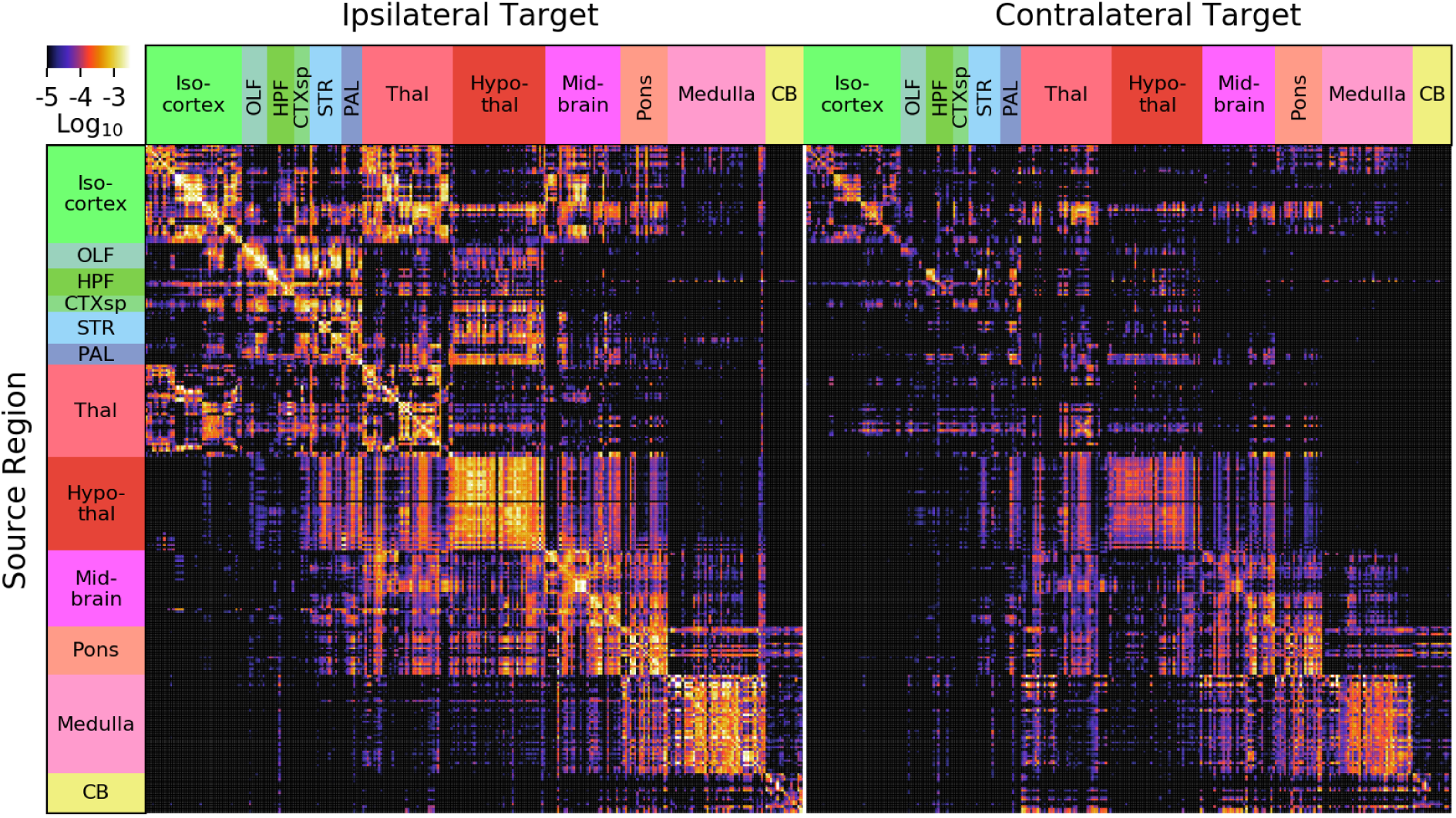
Whole-brain normalized connection density obtained from the regionalized voxel model. We show 292 regions divided into 12 major brain divisions. For visualization purposes, sources are shown on the rows and targets on the columns, the opposite convention as the mathematics in the text (*W*^*T*^ is pictured). The similarity between rows, e.g. in hypothalamus, is driven both by biological similarity and as a result of the model’s interpolation in the sources. The similarity between columns is purely biological, as the model does not interpolate in target space.

**Figure 3:**
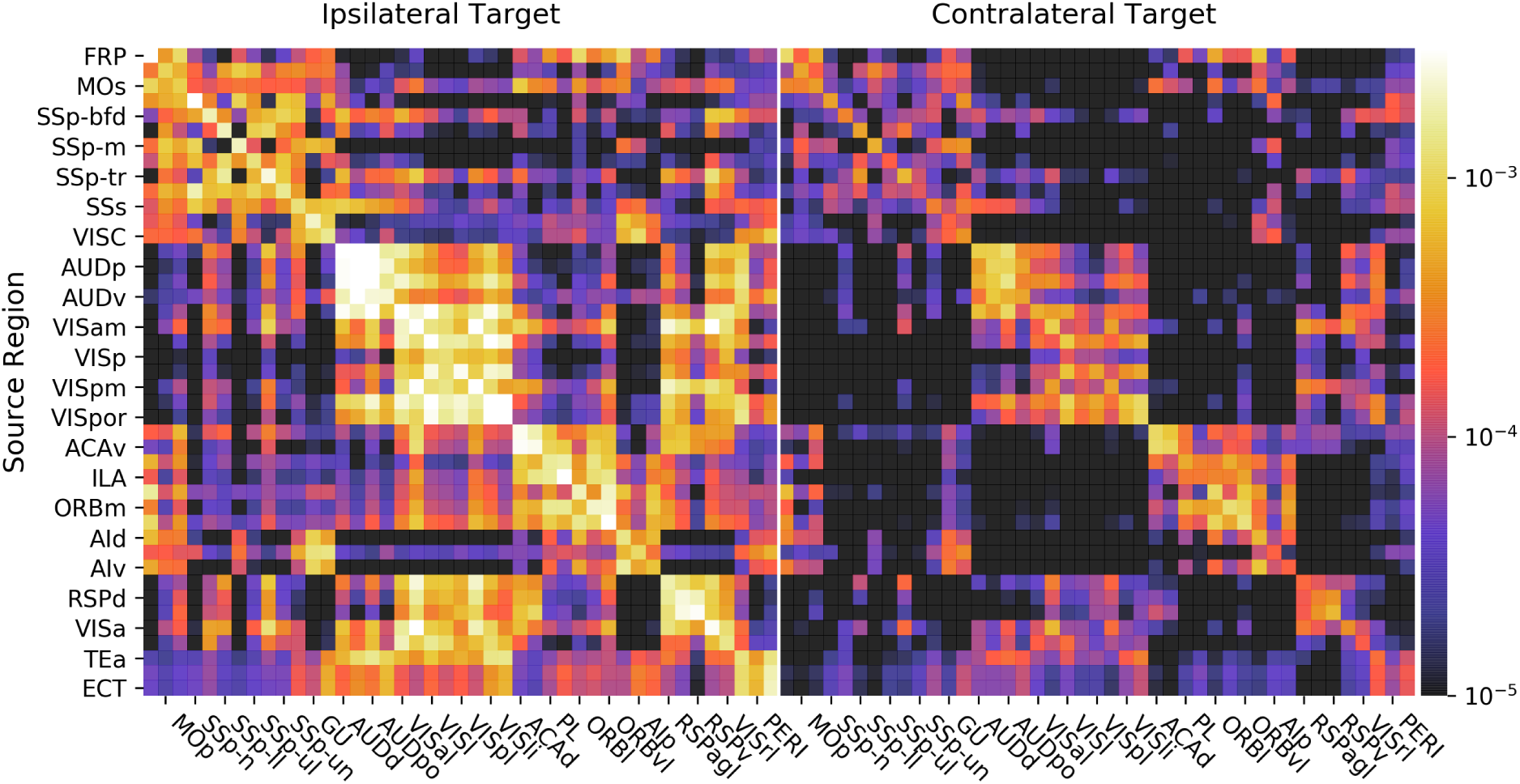
Isocortex normalized connection density from the regionalized voxel model. Again, we show sources as rows and targets as columns. The connectivity shows block structure, such as between the auditory and visual areas, as well as somato-motor areas. These blocks are functionally related areas connected into modules.

A number of features are evident in Figs. 2 and 3. First, there are patterns that arise from our smoothness assumption. The vertical banded structures (for example, the column near the right side of the Medulla structure in Figure 2) are due to smoothing in source but not target regions. In Fig. 3, note that the rows PERI, ECT, and TEa (the bottom three rows) and AUDv (upper middle) match closely. All four of these regions are very close to each other on the posterior, ventrolateral part of the cortex, so that smoothing causes them to be correlated. Also notice that, while the rows corresponding to these regions as sources are very similar, the columns corresponding are not nearly as much so. This highlights that our model interpolates only in the source, and not the target.

The second feature that is evident is the presence of blocks of strongly interconnected regions. These correspond to modules in the network, or regions that are more highly connected to each other than they are to the rest of the network. In Fig. 3, blocks of high connection density corresponding to somatomotor, auditory, visual, prefrontal, and medial areas can be visualized along the diagonal in the order listed. A systematic analysis of modularity revealed that the cortical network divides into 1–14 modules, depending on the choice of parameters (Harris et al., in submission).

In Table 1, we show the results of comparing homogeneous and voxel models. We fit the models using nested leave-one-out cross-validation. This allows us to evaluate both the voxel-scale and regionalized models’ error when predicting held-out data. We report mean squared error relative to the average squared norm of the prediction and data; see Eqn. 4. The model validation and training errors (goodness of fit, shown in parentheses) are reported at both voxel (Voxel MSE_rel_) and regional (Region MSE_rel_) levels. Additionally, we compare model performance using only the subset of experiments where every source region received repeated injections. More specifically, we only include experiments if the injection center of mass of at least 3 experiments are located in the same region. Without another injection, the only information about that region’s projections would come from our smoothness assumption interpolating nearby regions’ patterns. The computed relative error MSE_rel_ with this repeated dataset we call the “power to predict” (Region PTP).

**Table 1.** Table of cross-validated model errors, comparing both the voxel model and regionally homogeneous model. In each case the training error is in parentheses. Voxel MSE_rel_ refers to relative error, Eqn. 4, at the voxel level. This measure approximates the data normalized MSE for small errors, but is bounded to maximum of MSE_rel_ = 200%, which is achieved if either *Y* ^true^ or *Y* ^pred^ is zero and the other is not (see section 4.2.3). Region MSE_rel_ is the error found after regionalizing the voxel mode prediction. PTP (“power to predict”): Here, we evaluate MSE_rel_ for only those held out experiments where there was another injection in that region which was used for fitting.

| Superstructure | Model | Voxel $\text{MSE}_{\text{rel}}$ | | Region $\text{MSE}_{\text{rel}}$ | | Region PTP | |
| --- | --- | --- | --- | --- | --- | --- | --- |
| **Iscortex** | Voxel | 66% | (21%) | 33% | (11%) | 32% | (8%) |
|  | Homogeneous | - | - | 40% | (19%) | 31% | (17%) |
| **Olfactory Areas** | Voxel | 82% | (19%) | 41% | (7%) | 39% | (6%) |
|  | Homogeneous | - | - | 57% | (10%) | 31% | (9%) |
| **Cortical Subplate** | Voxel | 120% | (6%) | 104% | (2%) | 92% | (2%) |
|  | Homogeneous | - | - | 88% | (9%) | 72% | (9%) |
| **Hippocampus** | Voxel | 93% | (18%) | 56% | (17%) | 53% | (17%) |
|  | Homogeneous | - | - | 46% | (48%) | 42% | (48%) |
| **Striatum** | Voxel | 104% | (8%) | 46% | (2%) | 41% | (1%) |
|  | Homogeneous | - | - | 48% | (24%) | 48% | (25%) |
| **Pallidum** | Voxel | 103% | (8%) | 71% | (3%) | 45% | (5%) |
|  | Homogeneous | - | - | 68% | (4%) | 45% | (9%) |
| **Thalamus** | Voxel | 103% | (10%) | 59% | (9%) | 49% | (10%) |
|  | Homogeneous | - | - | 87% | (15%) | 86% | (23%) |
| **Hypothalamus** | Voxel | 59% | (58%) | 28% | (31%) | 21% | (8%) |
|  | Homogeneous | - | - | 41% | (5%) | 66% | (8%) |
| **Midbrain** | Voxel | 90% | (32%) | 30% | (13%) | 29% | (10%) |
|  | Homogeneous | - | - | 47% | (16%) | 43% | (17%) |
| **Pons** | Voxel | 99% | (11%) | 52% | (8%) | 51% | (8%) |
|  | Homogeneous | - | - | 69% | (31%) | 113% | (65%) |
| **Medulla** | Voxel | 94% | (39%) | 34% | (19%) | 35% | (20%) |
|  | Homogeneous | - | - | 68% | (45%) | 64% | (29%) |
| **Cerebellum** | Voxel | 189% | (8%) | 104% | (1%) | 95% | (6%) |
|  | Homogeneous | - | - | 81% | (26%) | 34% | (5%) |

From Table 1, we see that the relative training and validation errors are higher when evaluating error in the voxel space. This makes sense because this error captures mistakes we make in predicting spatial patterns of projections at the voxel, sub-regional level. That task is much more difficult than predicting regional patterns. The lowest voxel errors are in Isocortex, Olfactory Areas, and Hypothalamus. At the regional level we compare both the homogeneous model and the *regionalized* voxel model, which uses the voxel connectome to make a prediction that is then integrated across each region. At the regional level, our voxel model has lower regional validation errors than the homogeneous model in 8/12 major brain divisions. The training error is lower in 11/12 cases, since assuming smoothness is less biased than regional homogeneity. The regional PTP of the voxel model is lower than or roughly equal to (within 1%) the PTP of the homogeneous model in 8/12 cases. Results for training PTP are similar. In general, we find the highest errors (for either model) in major divisions which are small, composed of many small regions, and with large distance between injections. These statistics are summarized in Supplementary Table 2.

### 2.3 Visualizing the voxel-scale connectivity: cortical-cortico virtual injections

Visualization of our model faces two challenges: the matrix *W* contains *n* × *n* = *O*(10^11^) single precision floating point elements, and contains dense 3-D spatial structure. In order to address these challenges we opted to generate “artificial injections,” compute predicted projections from these injections, and visualize the resulting volumes. These artificial injections allow us to visualize the average projections from voxels of our choosing. This process is efficient, first because the matrix *W* is formed explicitly from *m* rank one components, so we only have to store *n* × *m* = *O*(10^8^) items. Standard tools, such as volume rendering and projection, can then be applied to visualize the model’s predictions.

In order to visualize model predictions in the Isocortex we make use of a curved cortical coordinate system. This coordinate system defines two dimensions over the surface of the cortex and one which is composed of steepest-descent paths from the pia surface to white matter. By projecting model predictions along these paths we can generate 2-D cortical projection maps which are faithful to the boundaries of Isocortical regions.

We display two such projections in Fig. 4. Here, we visualize the average over the columns of the matrix *W* corresponding to the projections from two major brain regions, marked by a red outline. We immediately observe strong ipsilateral projections to related areas. For instance, primary visual area VISp has a number of local projections to higher visual areas, Fig. 4a.

**Figure 4:**
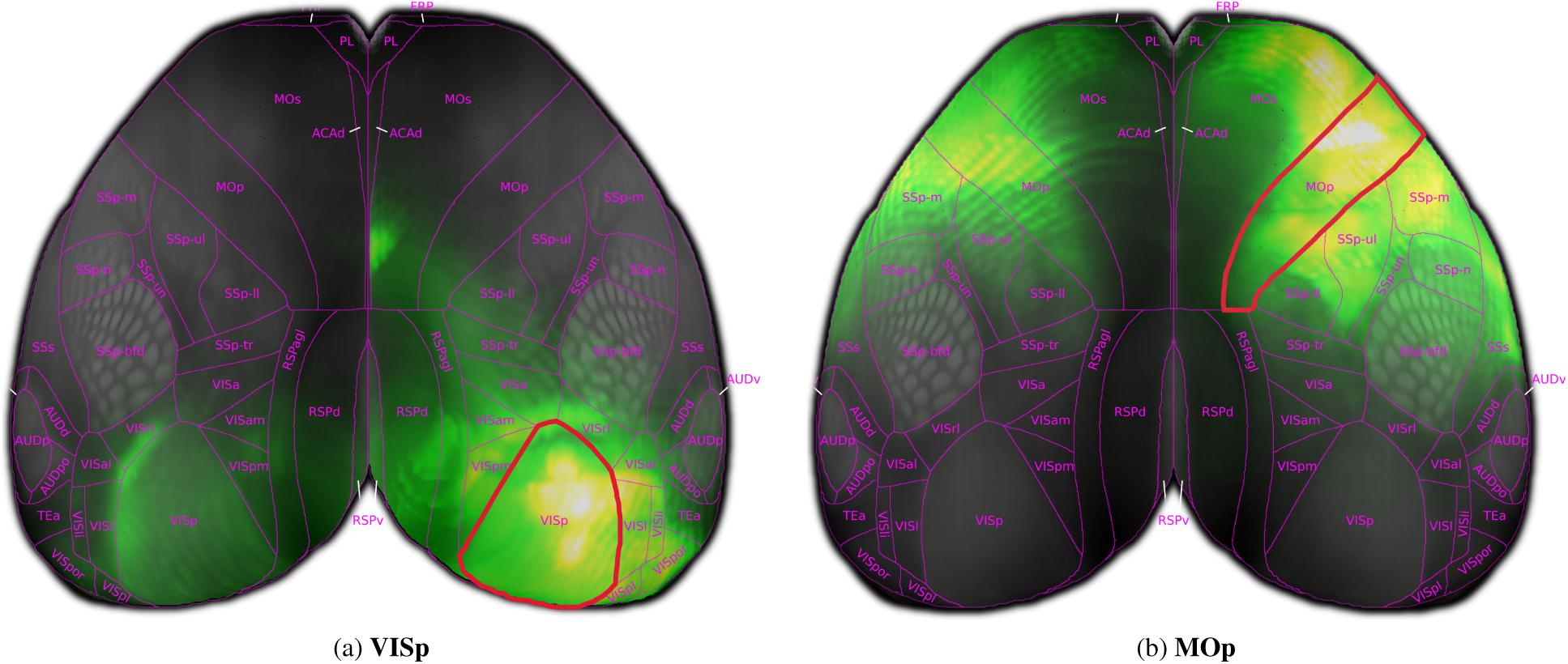
Model predicted cortical-cortico projections from virtual injections into the entire primary visual area (VISp) and primary motor area (MOp).

### 2.4 Weight distribution and its distance dependence

We compared multiple models for the distribution of weights: lognormal, inverse gamma, exponential and normal. We separately construct these models for ipsilateral and contralateral connections for the entire brain, and for connections within isocortex. For all these weight distributions, the best fit is for a lognormal distribution (using the Bayesian information Criteria, BIC; see Table 3). However, the results from the Kolmogorov-Smirnov test show that the fitted lognormal distributions fail to be statistically similar to the weights distributions for any of the divisions of the connections. Additionally, the logarithmically transformed weights fail to pass the Shapiro-Wilk test for normality at *α* = 0.05 level of significance. This is because the weights, depicted on the right hand side of Fig. 5, exhibit a skewed distribution.

**Figure 5:**
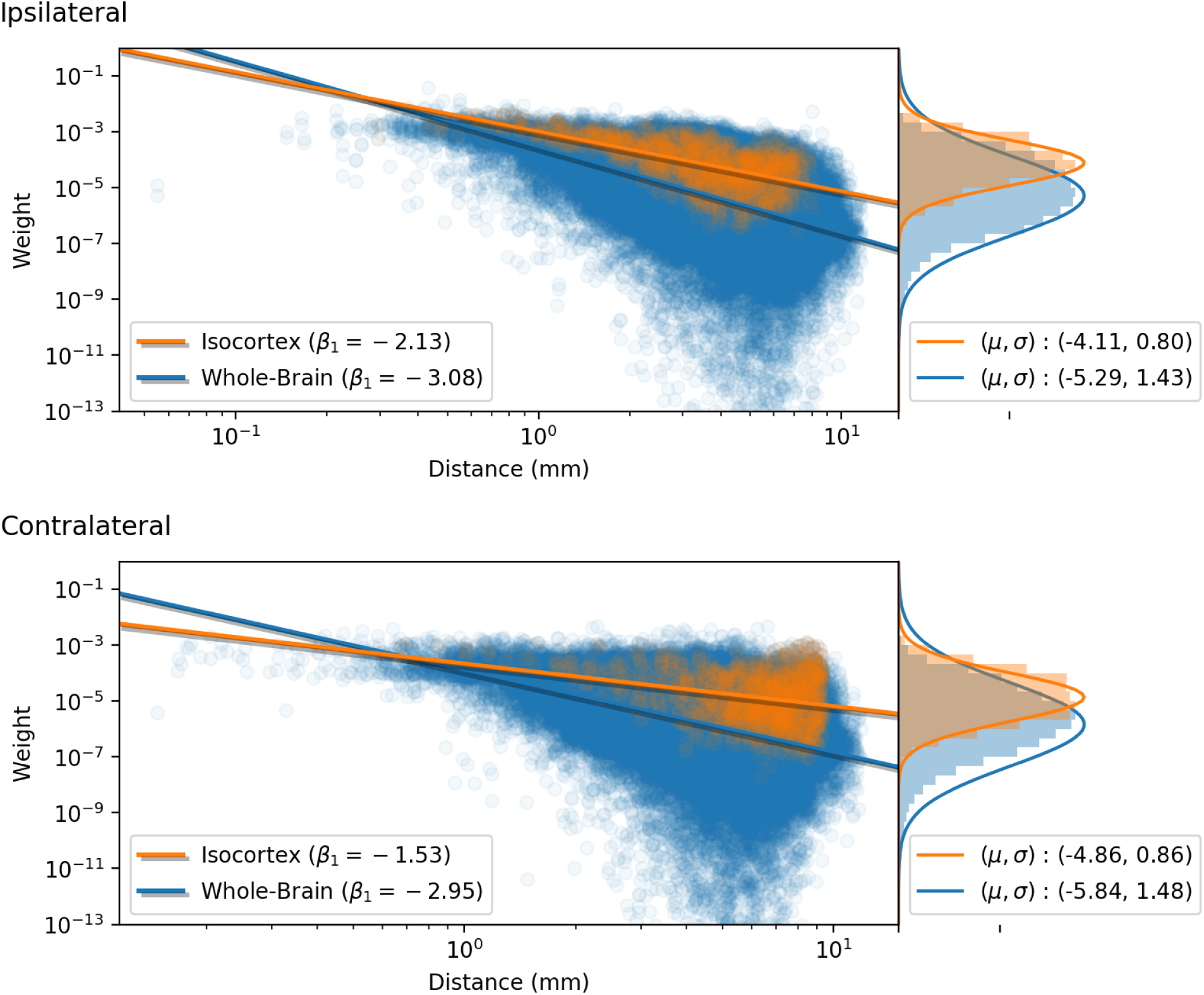
Normalized connection density produced by the regionalized voxel model (log scale) plotted against inter-region distance for 292 regions in the whole-brain (blue) and for only cortical-cortico connections (orange). The lines are linear least squares of the form log_10_(weight) = *β*_1_ log_10_(distance) + *β*_0_. The histograms on the right side show the distributions of weights as well as Gaussian fits of the log_10_(weight) distribution. Note that the standard deviations are slightly biased due to small weight outliers.

We have previously seen that a heterogeneous set of connections can be better fit by a mixture of lognormal distributions Oh et al. (2014). In a similar manner, we find the logarithmically transformed weights are best fit by a multiple component Gaussian mixture model (GMM) (see Table 4). The number of components was selected to minimize BIC resulting in a 5 component GMM for the whole brain, and 2-3 component GMM for cortical-cortico connections. With the exception of two components in each of the whole-brain mixture models, the components have similar valued weights, suggesting that different regions contribute to a non-homogeneous distribution of connection weights across the brain. However, it could also be the case that the empirical distribution of log-transformed weights is well-modeled by a unimodal distribution which is not Gaussian.

In Fig. 5, we show the dependence of extant connection weights on distance. We compared an exponential and a power-law fit. Using the Levenberg-Marquardt algorithm to fit nonlinear least squares problems, the root mean squared error (RMSE) was found to be slightly smaller for the power law fit.

### 2.5 Model performance compared with anatomical data

To evaluate how closely the model predictions aligned with experimental data, we compared the model predictions from both the homogeneous and regionalized voxel models with projection data for each of the 128 injections into the Isocortex in wild type mice from the Allen Mouse Brain Connectivity Atlas. For each injection experiment, we calculated the Pearson correlation between the normalized projection volume and the model prediction for both hemispheres in each of the 292 summary structures. Figure Fig. 6(a) shows the correlation values for each experiment grouped by source structure. The mean correlation coefficient for the regionalized Model was higher than for the homogeneous model (mean ± SD regionalized: 0.93 ± −0.08, homogeneous: 0.91 ± −0.12, p=0.03, paired t test), but each model outperformed the other for some sources.

**Figure 6:**
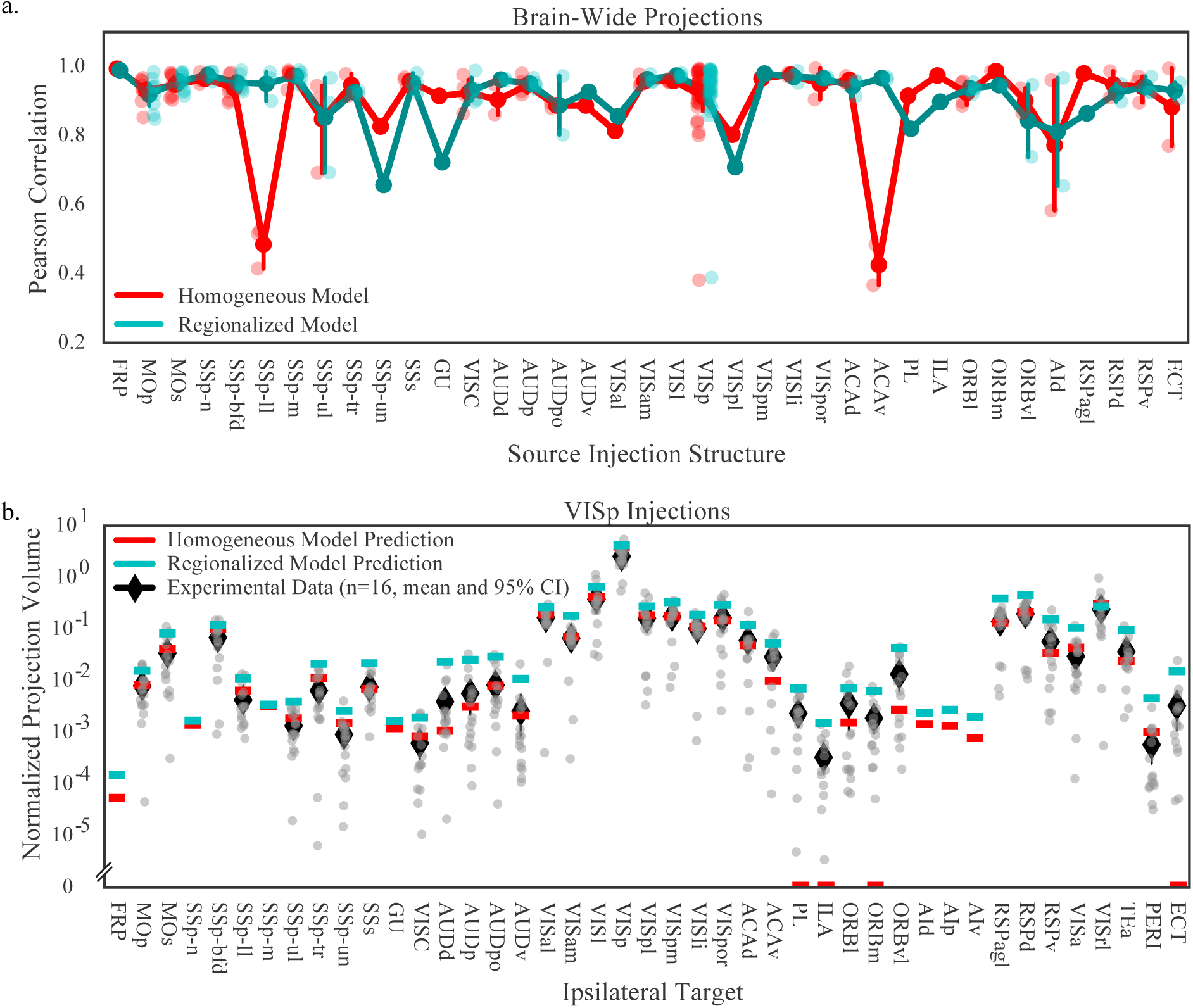
Model performance compared to human-curated “ground truth.” (a) Pearson correlation of model predictions and experimental data for injection experiments into 37 cortical source regions. Lighter colored points indicate correlation coefficients for individual experiments; darker points and lines indicate the mean and 95% confidence interval for all experiments in each source. (b) (log scale) Normalized projection volume in cortical targets from experimental data (gray) with corresponding predicted regional projection weights from the homogeneous model (red) and the regionalized voxel model (cyan) overlaid. Projection data from individual experiments is plotted as gray points and the mean and 95% confidence interval for each target are plotted as black diamonds (most confidence intervals are smaller than the markers). Normalized projection volume below 10^*−*6^ are truncated in this graph (5 data points).

To further explore the relationship between the model prediction and the experimental data, we also compared the predicted weights from both models with experimental data from a subset of experiments in which injections sites were *>* 95% contained within a single source region. There were 29 experiments that met this criterion; all were in the Isocortex with the following sources: AUDp (n=1), ENTl (n=1), MOp (n=2), MOs (n=1), RSPd (n=1), SSp-bfd (n=1), SSp-m (n=4), SSp-n (n=1), SSs (n=1), and VISp (n=16). Some of these experiments had been manually checked for segmentation errors, and where this data was available we multiplied the normalized projection volume by the manual call (1 for true positives, 0 for true negatives). For each of these ten sources, we plotted the normalized projection volume from the experimental data along with the predicted weights from both models for targets in the ipsilateral Isocortex.

Fig. 6(b) shows the plot for VISp, where we had a total of 16 experiments of which three were checked for true positive/true negatives. The weights predicted by the regional model were higher overall, but generally agreed with the experimental data as well as the homogeneous model predictions. The biggest difference between the two models was that the regionalized model correctly predicted weights for several targets that were missed by the homogeneous model (PL, ILA, ORBm, and ECT). All four of these targets were verified true positives but had a predicted weight of zero with the homogeneous model. On the other hand, both models predicted very small but non-zero weights for targets that were true negatives (FRP, SSp-n, AId, AIp, and AIv). Across all ten sources that were checked, the regionalized model routinely predicted weights for connections that were zero in the homogeneous model prediction, and overall there was a tendency for the regionalized model to predict higher weights, particularly for small structures.

Overall, we found the predictions of the regionalized voxel model to be more consistent than the homogeneous model, so that even when the predicted weight from the regionalized model was incorrect it was not off by a large margin. Conversely, the homogeneous model often performs well except when its predictions considerably differ from the experimental data. This is well-illustrated in Fig. 6(a), where the homogeneous model performs well overall, but very poorly for projections from SSp-ll and ACAv.

## 3 Discussion

In this study, we infer whole brain connectivity at a 100 µm voxel level from a set of brain-wide anterograde viral tracing experiments in young adult C57BL/6J mice (Oh et al. (2014), http://connectivity.brain-map.org/). The central assumption of the method is that brain-wide projections from nearby neurons within a brain region vary smoothly. Such a method, and its application to the visual system, has been described in Harris et al. (2016). However, using a smoothness prior simultaneously both at source level and at target level is computationally quite difficult, and led us to develop the simpler source-space interpolation presented here.

The tracing data which forms the basis of this study is based on anterograde viral tracers (Oh et al., 2014). Serial images are acquired with two photon tomography, at a sampling distance between planes of 100 µm. The in-plane resolution is much higher from the raw images (0.35 µm x 0.35 µm), but for the purpose of this study the data has been re-processed to an isotropic 100 µm resolution. The viral tracing methods used to generate the Allen Mouse Connectivity Atlas dataset results in two limitations affecting our ability to resolve connections. One limitation comes from the size of the injections, which have a typical radius of 0.3 mm (Table 2 provides the volume distribution through different brain regions). An even stronger limit comes from the typical distances between a voxel and the center of mass of an injection centroid being typically 0.5 mm (Table 2 reports this average in the “Inj. center distance” column). This distance is the consequence of the number of injections (429, of which we select 366 as described in Methods) being much smaller than the number of source voxels (2.5 × 10^5^). Thus the connections originating from a voxel have to be inferred from sources on average 0.5 mm away which average information over a radius of 0.3 mm Given the difference between source and target in accuracy of the data, applying the smoothness prior in the source space only seemed reasonable.

We compared the voxel and regional models’ ability to predict held out injection experiments. Although the errors for both are relatively high, the voxel model on average performs better. Furthermore, we are asking a lot of the models to predict such held out data when on average we only have a few injections per region.

We believe a good method to evaluate the model’s performance is to compare the predicted weights with a human-curated ground truth metric. We were able to make this comparison for a subset of injections well-contained in a few cortical sources. By comparing the models’ predictions with experimentally derived normalized projection volume values, we found two main differences between the regionalized model and the homogeneous model. Most importantly, the regionalized model predicts very weak but nonzero connections that the homogeneous model assigns zero weight. This is due to the inherent tendencies of the models to increase sparsity (homogeneous model) or to decrease to decrease sparsity (regionalized model). We verified some of the connections that were detected by the regional model and not the homogeneous model as true positives, but others were true negatives that were incorrectly assigned a weight by the regionalized model. The other main difference between the two models was in the prediction of weights for small target structures. Because the regionalized model is a linear smoother, it will tend to over predict weak connections for targets near regions with high connectivity. In choosing the appropriate model for an application, it is therefore important to consider the higher uncertainty of the presence of predicted weak connection.

We would like to emphasize that when analyzing this connectivity, especially from a graph theoretical perspective, one has also to be mindful of the correlations between connections originating from nearby sources that are introduced by the methods used. The spatial resolution of the connectivity is presented at 100 µm resolution: this is the native resolution for measurements of targets. However, at source level, the average distance to the closest injection is typically 0.5 mm (see Table 2), which limits the resolution. Also, many graph statistics may not be well-suited for studying such explicitly spatial graphs as ours.

Among multiple models for weight distributions, we found the log-normal being the best fit, in accordance with previous studies (e.g., Oh et al., 2014). Following this analysis, we found that a mixture of normal distributions better fit the distribution of log weights, which can be expected for heterogeneous neuronal populations. We analyzed the distance dependence of the connection weights and found that a power law dependence is a better fit than exponential. It is interesting that the power is close to −2 for the cortex (ipsilateral), which is primarily a 2-D structure, and close to −3 for the entire brain, which is 3-D. However, the weak scaling we observe only holds over roughly 1.5 orders of magnitude, so we prefer not to speculate too much about this result.

The voxel model enables quantitative characterization of the structural connectivity of the mouse brain. It is a significant improvement over the previously published homogeneous linear model (Oh et al., 2014), with tractable mathematics compared to the earlier voxel proposal (Harris et al., 2016). It offers improved predictions at region level, but, more importantly, it provides the connectivity at a much higher spatial resolution. This new model provides the necessary basis for studies of large-scale network structure, enabling discovery of general organizational rules for brain-wide systems which consist of both local and long-distance connections. A better understanding of these rules will lead to more accurate predictions of the directions of information flow, constrained by anatomy, and can be used by researchers interested in questions of structure-function relationships in the mouse brain.

## 4 Methods

### 4.1 Summary of data

The data were taken from 429 experiments using wildtype C57BL/6J mice. These data are available from the Allen Mouse Brain Connectivity Atlas at http://connectivity.brain-map.org/. We divide the brain into a set of *s* = 12 major brain divisions at a high level of the 3D Allen Mouse Brain Reference Atlas ontology. These major brain divisions are: Isocortex, Olfactory Bulb, Hippocampus, Cortical Subplate, Striatum, Pallidum, Thalamus, Hypothalamus, Midbrain, Pons, Medulla, Cerebellum. We also consider a finer partition (lower in the ontology) into a set of *r* = 292 regions. The major brain divisions form a disjoint partition of the brain, as do the regions. However, the regions are each contained within a given major brain division.

We conducted a hand-curation of the experiments to exclude those that have injections which substantially overlap multiple major brain divsions. For example, we removed experiments with large injection volumes spanning multiple subcortical major brain divisions and subcortical injections with substantial leakage of the tracer in the overlying cortex. Additionally, we removed four experiments having very little to no long-distance projections (small projection volume outside of the injection location). Overall, of the 429 experiments, we removed 63 experiments resulting in a total of 366 included experiments. We summarize these experiments used to fit our connectome in Table 2.

In our mathematical framework, the brain is a subset of ℝ^3^ which is discretized into a collection of *n* cubic voxels. Subsets of these voxels then correspond to the major brain division 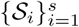 and regions 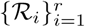. Each voxel *i* maps to a location in the brain, which we denote by *v*_*i*_ ∈ ℝ^3^. Each injection tracing experiment leads to an image of fluorescence throughout the brain. The fluorescence signal is reported as injection density (fraction of fluorescing pixels per voxel for voxels in the annotated injection site) and projection density (fraction of fluorescing pixels per voxel elsewhere). For the *e*th experiment, let *X*_:*,e*_ and *Y*_:*,e*_ denote the length *n* vectors of injection density and projection density, respectively. We also compute voxel coordinates of the center of mass of the injection density *c*_*e*_ ∈ ℝ^3^. For our estimator, we also compute the normalized projection density by normalized by the sum of the injection density, and denote this 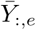. Note that 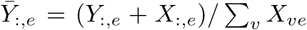, since we also include the injection pattern in the normalized projection density. Thus, the experimental data is this collection 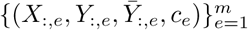 of length *n* vectors as well as the injection centers of mass for each experiment.

### 4.2 Multivariate nonparametric regression to infer voxel connectivity

We consider the problem of fitting a nonnegative, weighted adjacency matrix 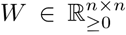 that is common across animals. Entry *W*_*ij*_ is the estimated projection density of neurons in voxel *j* to voxel *i*, if one unit of virus were delivered to voxel *j*. Each experiment consists of an injection *X*, and its projections *Y*, and we would like to find *W* so that so that *Y* ≈ *W X*. Uncovering the unknown *W* from multiple experiments (*X*_:*,e*_, *Y*_:*,e*_) for *e* = 1,…, *m* is then a multivariate regression problem. The unknown matrix *W* is a linear operator which takes images of the brain (injections) and returns images of the brain (projections).

Unlike the earlier work by Harris et al. (2016), we make two crucial simplifying approximations: First, we assume that in experiment *e* the injection is delivered to precisely one voxel, the injection center of mass *c*_*e*_. This removes the more difficult credit assignment problem of which voxels within each injection site contribute which projections. The method of Harris et al. (2016) solved this problem by essentially “dividing out” the injection correlations across experiments. Second, we assume that projections vary smoothly as we change the source voxel, i.e. the columns of *W* are smooth functions of the column voxel. However, we do not explicitly assume that the incoming projections to a target voxel vary smoothly as we move the target voxel, or smoothness in the rows. Smoothness in target space leads to dependencies among the output variables of the multivariate regression problem, making it a so-called *structured* regression problem, which are generally more difficult to solve. Note that, because the data tend to produce patterns of projections that are spatially smooth, and because we enforce smoothness in the source space, some target smoothness will naturally arise from the data and assumptions.

#### 4.2.1 Nadaraya-Watson connectome estimator

With these simplifying assumptions, we can now state the model. Our data are now the pairs of center of mass voxels *c_e_* and normalized injection densities *Y*_:*,e*_, which we assume arise from an injection of one unit of virus to the center of mass. Kernel regression is a standard non-parametric method for estimating a smooth univariate or multivariate function. For simplicity, we use the Nadaraya-Watson estimator (Nadaraya, 1964; Watson, 1964) to estimate the connectivity:

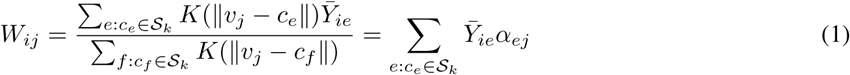

where

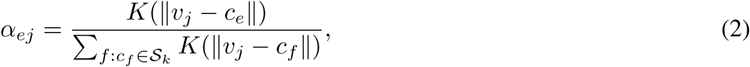

and *k* is the unique index such that 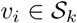, i.e. we only average over injections in the major brain division containing the source voxel. Furthermore, we can construct the matrices

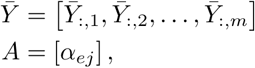

so that the connectome is written compactly as a rank *m* matrix 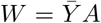, where 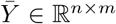 and *A* ∈ ℝ^*m*×*n*^. Note that each column in *A*, the coefficients *α*_*ej*_, has entries which sum to one.

The Nadaraya-Watson estimator, Eqn. (1), has a number of nice properties: It does not require any fitting, because the coefficients *α*_*ej*_ are given explicitly in terms of the center of masses and kernel, Eqn. (2). Furthermore, it forms a compressed rank *m* representation of *W* which is only as large as the data. However, it does suffer some drawbacks: It is well-known that the Nadaraya-Watson estimator is biased for data that is not sampled uniformly and near boundaries.

Note also that experiments with center of masses 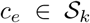 do not have any influence outside of 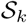. This is because we do not want to average over experiments in vastly different brain areas. Therefore, the coefficients *α*_*e,k*_ are decoupled across major brain divisions. Essentially, we fit a different model for each major brain division.

#### 4.2.2 Choice of spatial kernel

One needs to select a kernel to apply Eqn. (1). We use kernels of the polynomial family with finite support:

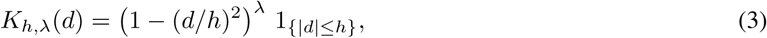

where *h* and *λ* are hyperparameters which set the size of the support and polynomial degree, respectively. The uniform kernel is in this family with *λ* = 0 and the Gaussian kernel is a limiting case as *λ* → ∞.

For polynomial kernels Eqn. (3), we have two hyperparameters to fit: the length scale *h* and exponent *λ*. However, we have a good lower bound for the length scale *h*, since we do not want any voxel in the brain to have zero connectivity. We therefore set *h* to be the maximum of the minimum distance from any voxel to the closest injection center of mass.

#### 4.2.3 Evaluating performance via cross-validation

To evaluate the performance of the model, we employ nested leave-one-out cross validation. In the inner loop, we fit *m* − 1 different models on sets of *m* − 2 experiments in order to perform model selection, wherein we fit the hyperparameters *h* and *λ* of the kernel function *K*. The best model is then evaluated against the held out experiment from the outer loop, and this process is repeated *m* different times. The performance metric we choose to use is mean square error relative to the average squared norm of the prediction and left out data:

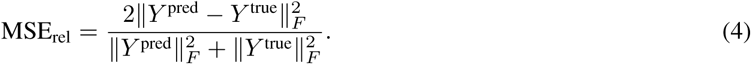

This choice of normalization prevents experiments with small ||*Y*|| from dominating the error, more heavily weighting experiments with larger signal (Harris et al., 2016).

The relative error in Eqn. (4) is approximately equal to the usual relative mean square error 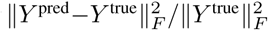 when *Y*^pred^ is close to *Y*^true^, and this is not too small. To see this, let *Y*^pred^ = *Y* ^true^ + *δ* where ||*δ*||_*F*_ ≤ ϵ and ||*Y*^true^||_*F*_ = *O*(1). Then, dropping the superscript “true” for clarity,

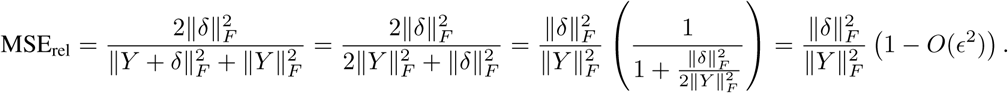

However, if *Y* is close to zero, our metric can be different. For example, if *Y* ^pred^ = 1 and *Y* ^true^ = 0.25, then MSE_rel_ = 2(1 − 0.25)^2^/(1^2^ + (0.25)^2^) = 106%. The usual relative mean square error would be (1 −0.25)^2^*/*(0.25)^2^ = 900%. If either *Y* ^true^ or *Y* ^pred^ is zero and the other is not, then MSE_rel_ = 200%, its maximum value.

Consider a set of experiments 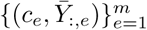, where *c*_*e*_ is the center of mass of the *e*th injection. Let *C* ∈ ℝ^*n*×*m*^ be the matrix of injection center indicators, with entries *C*_*ie*_ = 1_*{c*_*e*_ =*v*_*i*__}. Define the kernel matrix *A*^*c*^ ∈ ℝ^*m*×*m*^ as the kernel evaluated at the centers of mass, then this is just *A*_*c*_ = *AC*. Thus the model prediction of the center of mass projections is 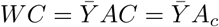.

We can perform leave-one-out cross validation efficiently after computing the coefficients *A*_*c*_ for a given set of data. If we leave out experiment *e*, the new model *W* ^(*−e*)^ predicts that the projections from *c*_*e*_ are 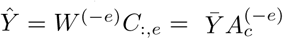, where 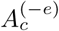 has the *e*th diagonal entry equal to zero and the corresponding column renormalized to sum to one. Therefore,

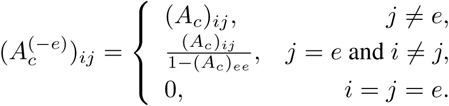

Extending the above result to compute the leave-one-out predictions for all of the experiments, we find that these are equal to 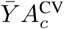, where

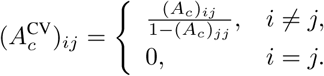

Thus, once we compute the coefficients *A*_*c*_, we set the diagonals equal to zero and renormalize the columns to obtain 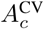. The leave one out cross-validation relative error of the voxel model is then

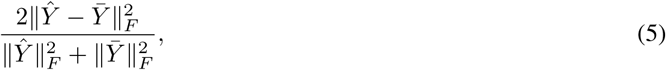

where 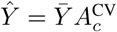 are the leave-one-out predictions.

### 4.3 Regionalized and homogeneous models

The application of Eqn. (1) results in a very large *n* × *n* voxel-scale connectivity matrix. Recall that we defined a parcellation of the brain into *r* regions 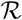. We would like to be able to compare this with extant regional connectomes, which are smaller *r* × *r* matrices. With this, we can define the regional projection matrix, Π;∈ ℝ^*r*×*n*^, with entries:

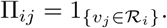

That is, the *i*th row of Π has ones in entries corresponding to voxels in region *i*. Therefore, for some vector *x* ∈ ℝ^*n*^ corresponding to a voxel image of the brain, the vector *x*^R^ = Π *x* has entries corresponding to the sum of *x* over regions. Furthermore, consider another matrix Π^*†*^ ∈ ℝ^n^×*r*, with entries:

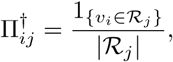

where 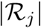 is the number of voxels in region *j*. Then Π^*†*^, operating from the left on a length *r* vector spreads the entries evenly over all of the voxels in a given region. Operating from the right, it averages over the voxels in a region. Note that Π Π^*†*^ = *I*_*r*_, so it is a right inverse of Π and in fact is a Moore-Penrose pseudoinverse.

With this notation, it becomes simple to convert voxel vectors and matrices into regional ones. We refer to the sum of the connection weights between two regions as the *connection strength* between the regions. Thus, the regional connection strength is given by

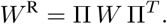

However, these regions may be vastly different sizes, in which case a measure normalized by source and/or target region size is more appropriate. We define the *normalized projection density* as the connection strength between two regions divided by the size of the source and target region. In this case, the matrix becomes

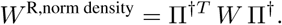

Finally, our last normalization only normalizes by the size of the source region, which we call *normalized connection strength*:

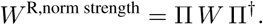

This normalization is necessary to compare directly with the homogeneous model.

#### 4.3.1 Fitting a homogeneous regional model

As in Oh et al. (2014), one could also fit a regional model where connection strengths are fixed across regions by working directly to data which are integrated over regions. We performed this for comparison and refer to the result as the *homogeneous* model. Let *X*^R^ = Π *X* and *Y* ^R^ = Π *Y*. Then the model fit to this regional data is found via nonnegative least squares as

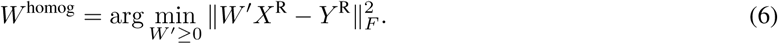

Note that the output *W* ^homog^ is a normalized connection strength, since entry (*i, j*) is the expected volume of fluorescence in region *i* per unit of virus in region *j*.

#### 4.3.2 Comparing the regionalized voxel model to the homogeneous model

Let 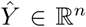 be the voxel prediction, we can compute the regionalized prediction 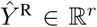 by projecting the voxel predictions into the regional space: 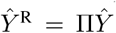. It is important to note that although the comparison between the regionalized voxel model and the homogeneous model are done in the same space ℝ^*r*^, the predictions themselves are slightly different. The regionalized voxel predictions 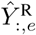 are the predicted result of a unit injection into the center of mass of the injection *c*_*e*_, whereas the regional prediction 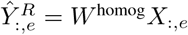 is the regional prediction of the projection from the full injection *X*_:*,e*_.

## 5 Supplementary materials

All of the data used to construct these models is available at http://connectivity.brain-map.org. The curved coordinate system is available at http://help.brain-map.org/download/attachments/2818171/ Mouse Common Coordinate Framework.pdf. All code used to build this model will be made available at the time of publication from https://github.com/AllenInstitute/high_resolution_data-driven_model_of_the_mouse_connectome.

## 6 Acknowledgements

This work was supported by the Allen Institute for Brain Science. The authors wish to thank the Allen Institute founders, Paul G. Allen and Jody Allen, for their vision, encouragement and support. Additionally, the project described was supported in part by the National Institute of Aging Award Number R01AG047589 to J.A.H. Its contents are solely the responsibility of the authors and does not necessarily represent the official views of the National Institutes of Health. K.D.H. was supported by the Big Data for Genomics and Neuroscience NIH training grant. K.D.H. and E.S.-B. were supported by NSF DMS grants 1122106 and 1514743.

## 7 Author Contributions

Conceptualization S.M., E.S.-B., K.D.H.; Methodology, Formal Analysis, Investigation J.E.K., K.D.H.; Validation J.E.K., K.D.H., J.D.W., J.A.H.; Software J.E.K., K.D.H., N.G.; Data Curation J.A.H.; Visualization J.E.K., K.D.H., J.D.W.; Writing J.E.K., K.D.H., J.D.W., S.M., with review and editing from N.G., J.A.H., E.S.-B.; Supervision H.Z., J.A.H., E.S.-B., S.M.

## 8 Supplement

### 8.1 Technical Terms

- **Connection Strength:** The sum of the connection weights from all voxels in a source region 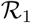 to all voxels in a target region 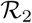, denoted 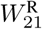.
- **Edge Density:** The total number of edges in a graph over the total possible number of edges in a graph. In our case, since our graphs are directed, it is 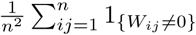.
- **Frobenius Norm:** The 2-norm of a matrix viewed as a vector: 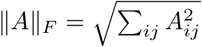
- **Normalized Connection Density:** The connection strength from 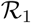 to 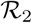 divided by the product of their sizes: 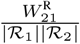
- **Normalized Connection Strength:** The connection strength from 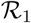 to 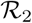 divided by the size of the source region: 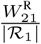.
- **Radial Basis Function:** A monotonically decreasing, nonnegative function.
- **Regions:** The set of 292 brain structures (also called ‘summary structures’) from the 3D Allen Mouse Brain Reference Atlas.
- **Major Brain Divisions:** The set of 12 major brain divisions (also called ‘coarse’ structures) from the 3D Allen Mouse Brain Reference Atlas: Isocortex, Olfactory Areas, Hippocampus, Cortical Subplate, Striatum, Pallidum, Thalamus, Hypothalamus, Midbrain, Pons, Medulla, Cerebellum.
- **Voxel:** A 3-D cubic volume element; the generalization of a pixel.
- **Wildtype mouse (C57BL/6J):** Mice of strain C57BL/6J which have not been genetically altered.

### 8.2 Supplemental Tables

**Table 2:** Summary statistics for included data.

| Major Brain Division | Expts. | Regions | Multiple expt. regions | Total Vol. (mm<sup>3</sup>) | Mean Region Vol. (mm<sup>3</sup>) | Mean Inj. Vol. (mm<sup>3</sup>) | Inj. center distance (mm) |
| --- | --- | --- | --- | --- | --- | --- | --- |
| Isocortex | 126 | 43 | 37 | 61.88 | 1.44 | 0.23 | 0.51 |
| Olfactory Areas | 21 | 11 | 7 | 23.28 | 1.89 | 0.12 | 0.71 |
| Hippocampus | 43 | 12 | 9 | 21.35 | 1.68 | 0.12 | 0.51 |
| Cortical Subplate | 8 | 7 | 2 | 4.43 | 0.61 | 0.15 | 0.88 |
| Striatum | 26 | 14 | 7 | 22.61 | 1.52 | 0.16 | 0.67 |
| Palidum | 10 | 9 | 2 | 4.75 | 0.47 | 0.12 | 0.61 |
| Thalamus | 43 | 40 | 25 | 10.30 | 0.24 | 0.11 | 0.40 |
| Hypothalamus | 34 | 41 | 14 | 7.64 | 0.15 | 0.33 | 0.41 |
| Midbrain | 43 | 33 | 12 | 18.66 | 0.46 | 0.21 | 0.46 |
| Pons | 16 | 21 | 7 | 8.51 | 0.30 | 0.15 | 0.60 |
| Medulla | 39 | 43 | 18 | 15.74 | 0.30 | 0.23 | 0.49 |
| Cerebellum | 19 | 17 | 7 | 27.22 | 1.57 | 0.08 | 0.76 |

**Table 3:** Model selection for the best fit distribution for the connectivity weights. The best-fit model was chosen through minimizing Bayes Information Criteria (BIC). BIC is based on the liklihood function with an additional penalty for the number of model parameters. The weight distributions of both ipsilateral and contralateral whole-brain and cortical-cortico connection weights were best fit by a lognormal distribution. However, these fits were not significantly similar to any of the above weights distributions as determined by the Kolmogorov-Smirnov test at a *α* = 0.5 level of significance as seen by the above p-values.

| Division | Projection Hemisphere | BIC |  |  |  |  |
| --- | --- | --- | --- | --- | --- | --- |
|  |  | Lognormal |  | Inverse Gamma | Exponential | Normal |
| Whole-Brain | Ipsilateral | -1.61e6 | $p = 2e-111$ | -1.39e6 | -1.09e6 | 8.30e9 |
| | Contralateral | -1.79e6 | $p = 2e-88$ | -1.58e6 | -1.29e6 | 3.37e11 |
| Isocortex | Ipsilateral | -2.70e4 | $p = 0.02$ | -2.62e4 | -2.57e4 | -2.17e4 |
| | Contralateral | -3.36e4 | $p = 0.01$ | -3.29e4 | -3.18e4 | -2.74e4 |

**Table 4:** Fitted gaussian mixture model parameters for the whole-brain and cortical-cortico logrithmically scaled connectivity weight distributions broken down by projection hemisphere. In both cases, the logrithmically transformed weights distributions failed to pass the Shapiro-Wilk test for normality, so we fit a mixture of gaussians to each of the logrithmically transformed distributions. As in Table 3, the number of gaussian components was selected to minimize the Bayesian Information Criteria (BIC). With the exception of two of the whole-brain components, the individual weights of the fitted gaussians are similar, possibly resulting from different regions contribute to a non-homogeneous distribution of connection weights across the brain.

| Division | Projection Hemisphere | Component | $\mu$ | $\sigma$ | weight |
| --- | --- | --- | --- | --- | --- |
| Whole-Brain | Ipsilateral | 1 | -6.21 | 0.24 | 0.29 |
|  |  | 2 | -5.02 | 0.21 | 0.33 |
|  |  | 3 | -11.37 | 9.71 | 0.01 |
|  |  | 4 | -7.56 | 0.66 | 0.08 |
|  |  | 5 | -3.81 | 0.26 | 0.29 |
|  | Contralateral | 1 | -4.34 | 0.26 | 0.26 |
|  |  | 2 | -8.18 | 0.84 | 0.07 |
|  |  | 3 | -6.71 | 0.26 | 0.29 |
|  |  | 4 | -5.53 | 0.19 | 0.36 |
|  |  | 5 | -12.36 | 13.07 | 0.01 |
| Isocortex | Ipsilateral | 1 | -3.23 | 0.15 | 0.35 |
|  |  | 2 | -4.21 | 0.13 | 0.44 |
|  |  | 3 | -5.19 | 0.22 | 0.21 |
|  | Contralateral | 1 | -5.52 | 0.32 | 0.50 |
|  |  | 2 | -4.20 | 0.29 | 0.50 |

**Table 5:** Model comparison of the best fit relation between normalized connection density and pairwise distances between anatomical regions. The power law and exponential relations were fit through a nonlinear optimization algorithm (Levenberg-Marquadt) to minimize the mean squared residuals. The performance of the two models is similar, with the powerlaw relation having slightly lower root mean squared error (RMSE).

| Division | Projection Hemisphere | RMSE |  |
| --- | --- | --- | --- |
|  |  | Power Law | Exponential |
| Whole-Brain | Ipsilateral | 2.85 | 2.89 |
|  | Contralateral | 3.14 | 3.19 |
| Isocortex | Ipsilateral | 1.40 | 1.48 |
|  | Contralateral | 1.89 | 1.92 |

## References

Bock, D. D., Lee, W.-C. A., Kerlin, A. M., Andermann, M. L., Hood, G., Wetzel, A. W.,… Reid, R. C. (2011, March). Network anatomy and in vivo physiology of visual cortical neurons. Nature, 471(7337), 177–182. doi: 10.1038/nature09802

Bota, M., Dong, H.-W., & Swanson, L. W. (2003, August). From gene networks to brain networks. Nature Neuroscience, 6(8), 795–799. doi: 10.1038/nn1096

Felleman, D. J., & Van Essen, D. C. (1991, January). Distributed Hierarchical Processing in the Primate. Cerebral Cortex, 1(1), 1–47. doi: 10.1093/cercor/1.1.1

Gămănuţ, R., Kennedy, H., Toroczkai, Z., Ercsey-Ravasz, M., Van Essen, D. C., Knoblauch, K., & Burkhalter, A. (2018, February). The Mouse Cortical Connectome, Characterized by an Ultra-Dense Cortical Graph, Maintains Specificity by Distinct Connectivity Profiles. Neuron, 97(3), 698–715.e10. doi: 10.1016/j.neuron.2017.12.037

Glickfeld, L. L., Andermann, M. L., Bonin, V., & Reid, R. C. (2013, February). Cortico-cortical projections in mouse visual cortex are functionally target specific. Nature Neuroscience, 16(2), 219–226. doi: 10.1038/nn.3300

Harris, K. D., Mihalas, S., & Shea-Brown, E. (2016). High resolution neural connectivity from incomplete tracing data using nonnegative spline regression. In Neural Information Processing Systems.

Jenett, A., Rubin, G. M., Ngo, T.-T. B., Shepherd, D., Murphy, C., Dionne, H.,… Zugates, C. T. (2012, October). A GAL4-Driver Line Resource for Drosophila Neurobiology. Cell Reports, 2(4), 991–1001. doi: 10.1016/j.celrep.2012.09.011

Kleinfeld, D., Bharioke, A., Blinder, P., Bock, D. D., Briggman, K. L., Chklovskii, D. B.,… Sakmann, B. (2011, September). Large-Scale Automated Histology in the Pursuit of Connectomes. The Journal of Neuroscience, 31(45), 16125–16138. doi: 10.1523/JNEUROSCI.4077-11.2011

Kuan, L., Li, Y., Lau, C., Feng, D., Bernard, A., Sunkin, S. M.,… Ng, L. (2015, February). Neuroinformatics of the Allen Mouse Brain Connectivity Atlas. Methods, 73, 4–17. doi: 10.1016/j.ymeth.2014.12.013

Laramée, M.-E., & Boire, D. (2015). Visual cortical areas of the mouse: Comparison of parcellation and network structure with primates. Frontiers in Neural Circuits, 8, 149. doi: 10.3389/fncir.2014.00149

Majka, P., Chaplin, Tristan A., Yu, Hsin-Hao, Tolpygo, Alexander, Mitra, Partha P., Wójcik, Daniel K., & Rosa, Marcello G.P. (2016, April). Towards a comprehensive atlas of cortical connections in a primate brain: Mapping tracer injection studies of the common marmoset into a reference digital template. Journal of Comparative Neurology, 524(11), 2161–2181. doi: 10.1002/cne.24023

Nadaraya, E. A. (1964). On Estimating Regression. Theory of Probability and its Applications, 9(1), 141–2.

Oh, S. W., Harris, J. A., Ng, L., Winslow, B., Cain, N., Mihalas, S.,… Zeng, H. (2014, April). A mesoscale connectome of the mouse brain. Nature, 508(7495), 207–214. doi: 10.1038/nature13186

Peng, H., Tang, J., Xiao, H., Bria, A., Zhou, J., Butler, V.,… Long, F. (2014, July). Virtual finger boosts three-dimensional imaging and microsurgery as well as terabyte volume image visualization and analysis. Nature Communications, 5. doi: 10.1038/ncomms5342

Sethi, S. S., Zerbi, V., Wenderoth, N., Fornito, A., & Fulcher, B. D. (2017). Structural connectome topology relates to regional BOLD signal dynamics in the mouse brain. Chaos: An Interdisciplinary Journal of Nonlinear Science, 27(4), 047405. doi: 10.1063/1.4979281

Sporns, O. (2010). Networks of the Brain (1st ed.). The MIT Press.

Stafford, J. M., Jarrett, B. R., Miranda-Dominguez, O., Mills, B. D., Cain, N., Mihalas, S.,… others (2014). Large-scale topology and the default mode network in the mouse connectome. Proceedings of the National Academy of Sciences, 111(52), 18745–18750.

Wang, X.-J., & Kennedy, H. (2016). Brain structure and dynamics across scales: In search of rules. Current Opinion in Neurobiology, 37(Supplement C), 92–98. doi: https://doi.org/10.1016/j.conb.2015.12.010

Watson, G. S. (1964). Smooth Regression Analysis. Sankhyā: The Indian Journal of Statistics, Series A (1961-2002), 26(4), 359–372.

White, J. G., Southgate, E., Thomson, J. N., & Brenner, S. (1986, November). The Structure of the Nervous System of the Nematode Caenorhabditis elegans. Philosophical Transactions of the Royal Society of London B: Biological Sciences, 314(1165), 1–340. doi: 10.1098/rstb.1986.0056

